# StrainXpress: strain aware metagenome assembly from short reads

**DOI:** 10.1101/2022.03.23.485539

**Authors:** Xiongbin Kang, Xiao Luo, Alexander Schönhuth

## Abstract

Next-generation sequencing based metagenomics has enabled to identify microorganisms in characteristic habitats without the need for lengthy cultivation. Importantly, clinically relevant phenomena such as resistance to medication, virulence or interactions with the environment can vary already within species. Therefore, a major current challenge is to reconstruct individual genomes from the sequencing reads at the level of strains, and not just the level of species. However, strains of one species can differ only by minor amounts of variants, which makes it difficult to distinguish them. Despite considerable recent progress, related approaches have remained fragmentary so far. Here, we present StrainXpress, as a comprehensive solution to the problem of strain aware metagenome assembly from next-generation sequencing reads. In experiments, StrainXpress reconstructs strain-specific genomes from metagenomes that involve up to more than 1000 strains, and proves to successfully deal with poorly covered strains. The amount of reconstructed strain-specific sequence exceeds that of the current state-of-the-art approaches by on average 26.75% across all data sets (first quartile: 18.51%, median: 26.60%, third quartile: 35.05%).

## Introduction

Metagenomics reveal the composition of complex microbial communities. Therefore, metagenomics facilitate to study the interactions and the environmental impact of microorganisms within their communities. In particular, next generation sequencing (NGS) has meant a major boost for metagenomics. The decisive advantages of NGS are to analyze the DNA isolated from environments of interest at little expense, and without the need to culture samples [21]. In the past decade NGS has been successfully applied to explore microbial communities from soil [10], ocean [25], human [23], and environments exposed to extreme conditions [35], among others.

A current major challenge is to assemble the individual genomes that make part of the metagenome. Unlike reference assisted assemblies, *de novo* assemblies are free of biases and can highlight individual genomes for which applicable reference genomes are still missing. Spurred by the great general interest, various *de novo* metagenome assembly methods have been developed. A non-exhaustive selection of state-of-the-art approaches encompasses IDBA-UD [28], SPAdes [5], Minia3 [8] and MEGAHIT [18].

The currently leading approaches successfully identify and reconstruct individual genomes at the level of species. However, their abilities in terms of distinguishing genomes at the level of strains leaves substantial room for improvements. Importantly, although belonging to the same species, strains can exhibit significant differences in terms of their interactions and the impact within the environment they belong to [37]. Moreover, strains can differ in terms of clinically relevant phenomena: for example, while various strains of *E. coli* are harmless, produce vitamin K [36] or suppress pathogenic bacteria [15], other *E. coli* strains can cause serious inflammatory processes or just poison food, such as *E. coli* EC958 [34] or *E. coli* O157:H7 [16]. Since the primary purpose of metagenomics is to analyze the interactions of the genomes with each other, and their impact on the environment they are drawn from, identifying genomes at the level of strains can be imperative.

Here and in the following, a “strain” is defined to be a viral or bacterial haplotype, as a contiguous genomic sequence that is supported by sufficiently abundant amounts of sequencing reads. We do this in agreement with related work on the topic [38], while we are aware of the possibly different meaning of a “strain” in terms of classic taxonomy or other concepts relevant in the evolution of microorganisms.

When investigating the shortcomings of prior approaches, one realizes that all of them follow the de Bruijn graph (DBG) assembly paradigm. Among other things, this implies to chop sequencing reads into smaller pieces of equal length k (“k-mers”). One then tries to identify paths in the DBG, defined by nodes reflecting k-mers, and edges reflecting overlaps of length k-1. So, in DBG based assembly, one trades off sequence length for benefits that usually relate to computational efficiency, which is supported by the efficient data structures that DBGs give rise to [30]. The trade-off is generally justified by the huge volume of the majority of contemporary NGS data sets [20].

Chopping reads into smaller pieces induces an obvious loss of information, however: the genetic linkage of sequential variants at distance greater than k can no longer be tracked [9]. Without additional precautions, which usually involve to consider reads at their full length in a post-hoc analysis, the cutting of reads into pieces of length k can severely hamper to distinguish between genomes of high identity^1^: as soon as the variants that are characteristic of the genomes are separated by more than k positions within their genomes, genetic linkage of variants can no longer be observed.

Instead, considering reads at their full length optimally preserves information about co-occurring (i.e. linked) mutations that are characteristic for the strains. This explains why approaches that do not require to chop reads into smaller pieces have recently gained considerable attention. The predominant data structure that supports such approaches are overlap graphs (OGs). Unlike DBGs, OGs do not require to work with k-mers, and have been the traditional counterpart of DBGs. First of all, however, OG based approaches traditionally require considerably more computational resources, which highlights the popularity of DBG based approaches yet another time.

Nonetheless, if one can get OG based approaches to run in an affordable amount of runtime and memory, OG based approaches do have clear advantages over DBG based approaches, because OG based approaches optimally preserve the identity of the haplotypes within the mix of genomes. This situation sets the framework for the currently driving methodical challenges in metagenome assembly: overcoming the computational bottlenecks of OG based approaches is key to success when aiming at considerably improved haplotype/strain aware assemblies of metagenomes.

The computational bottlenecks of OG based approaches are manifold. Recent works have demonstrated how to overcome single such bottlenecks, one at a time: first, an OG based solution for strain aware metagenome *gene* assembly (which addresses to reconstruct only the sequences of the genes within the genomes) was presented [13]. Subsequently, an OG based solution for the viral quasispecies assembly problem was suggested [2]. The approach specializes in virus genomes, which means that genomes are only of short length, whereas the coverage of the individual genomes is large. Subsequently, it was shown how to construct OGs for genomes spanning several millions of nucleotides, which enabled to assemble genomes of higher, but *fixed ploidy* in a ploidy-aware manner [3]. Further, an OG based approach was suggested that clusters raw metagenome sequencing reads into species specific groups without the need for a reference genome [4]. The motivation for this study was the fact that strain aware assembly tools could conveniently pick up clusters, while they failed to produce good assemblies when dealing with the full data set [4]. Beyond these OG based approaches that address to work with short NGS reads, it may be important to realize that OGs are the predominant data structure when assembling the genomes of vertebrates in a ploidy aware manner using third-generation sequencing reads [7, 17], which provides general motivation for pursuing OG based approaches also in metagenomics.

Overall, recent progress on OG based haplotype / strain aware assembly of genomes from mixed samples has been very promising on the one hand, but has remained fragmentary on the other hand. However, a comprehensive tool that synthesizes the recent progress by seamlessly combining the fragmentary work presented so far, and by modifying and adding whatever needs to be modified and added has not been presented so far.

The goal of this paper is to establish such a comprehensive solution and provide easy-to-use software that implements it. In this, this study suggests a considerable improvement over the earlier fragmentary, proof-of-concept approaches presented so far. Beyond comprehensiveness and easy usage an additional goal is provide an approach that is sufficiently lightweight; we recall that, if implemented in a naive manner, OG based approaches tend to be computationally (overly) demanding. Of course, however, the central purpose of our approach is to deliver strain-aware metagenome assemblies of utmost quality, and, so, to establish a considerable step up in NGS based

## Materials and Methods

### Overview

We present StrainXpress, as a tool that realizes all goals formulated towards the end of the Introduction: to the best of our knowledge, StrainXpress is the first comprehensive OG based approach by which to compute strain aware assemblies of metagenomes from NGS (Illumina type) reads. In the following, we will outline the methodical basis of the approach. In Results, we will demonstrate the superiority of StrainXpress with respect to the most relevant aspects. As a brief summary of its achievements, StrainXpress appears to be the only approach so far to deliver *strain-resolved* assemblies of metagenomes from NGS reads.

We recall that StrainXpress is based on overlap graphs (OGs) in the majority of its algorithmic routines. In the following, we first discuss the workflow of StrainXpress, as a high-level description of its algorithmic approach.

Subsequently, we will provide the full range of methodical details for each of the steps. We will also provide descriptions of the simulated and real data sets used in our experiments, and define the criteria by which we evaluate the assemblies.

### Workflow

From a general perspective, StrainXpress pursues a divide and conquer strategy that combines partial improvements into a comprehensive solution: while dividing refers to clustering reads into smaller portions and assembling strain-specific contigs for each cluster, conquering refers to collecting all contigs from each cluster, and assembling them further into longer strain-specific genomes in a global manner. Importantly, all of these steps—clustering, cluster based assembly of (strain-specific) contigs and global assembly of cluster derived contigs—are OG based.

See Figure 1 for an illustration. As just pointed out, the workflow consists of three steps. We recall that the first two steps reflect the divide stage, while the third step reflects the conquer stage:

**Figure 1.**
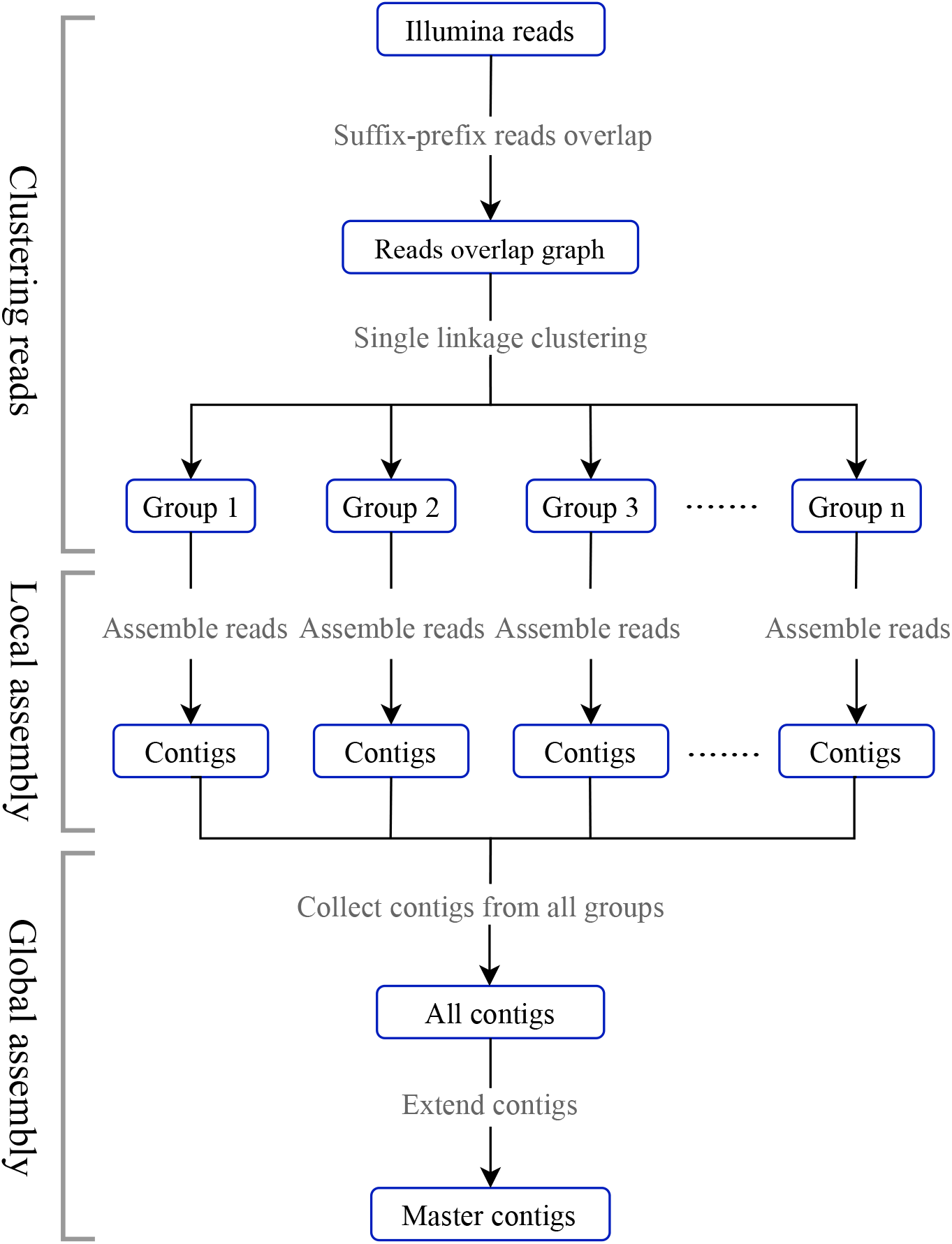
Workflow of StrainXpress. StrainXpress consists of 3 stages, ‘Clustering Reads’ (1), ‘Local Assembly’ (2), and ‘Global Assembly’ (3). All stages are based on overlap graphs as underlying data structure. The workflow follows a ‘Divide-And-Conquer’ strategy. While (1) and (2) reflect the ‘Divide’ part, (3) reflects the ‘Conquer’ part.

1. *Clustering reads* determines relatively small groups of reads that are likely to stem from identical species.
2. *Local assembly* assembles reads within (species) clusters into strain-aware contigs using cluster-specific overlap graphs. Strain-aware assembly is computationally feasible, because clusters are both sufficiently small and biologically coherent.
3. *Global assembly* takes in cluster-specific, strain-aware contigs from (2) and extends them by connecting them across clusters. This stage is based on a “master” overlap graph as underlying data structure, where nodes refer to cluster-specific contigs, and edges indicate sufficient and coherent overlap. The resulting “master contigs” are the output of StrainXpress.

As above-mentioned, StrainXpress draws inspiration from the (so far fragmentary) OG based work that was presented earlier. In particular, it was described how to cluster the reads of metagenomes into species-specific clusters [4, OGRE] based on a hierarchical single-linkage clustering strategy. Here, similar in spirit, we cluster NGS reads also following a single-linkage protocol. However, here we avoid the computationally expensive machine learning (ML) based routine by which to evaluate read overlaps [4], and replace it with an algorithmic protocol that is computationally inexpensive. We find that the substantial improvements in terms of speed offset the negligible losses in terms of overlap quality. Importantly, note that overlap quality and species consistency of clusters is not a primary goal here, which differs from the objectives formulated in the earlier study [4].

For step (2), we adopt an OG based, ploidy aware assembly strategy suggested earlier [3, POLYTE]. Here, it serves as a generic template for the conquer stage of StrainXpress. The decisive adaptation is to replace the FM index based computation of approximate overlaps as originally proposed [3] with minimizer bases schemes, as implemented by Minimap2 [19]. For that, the key insight has been to realize that Minimap2, although not primarily meant to deal with short reads, achieves improvements over the FM index based overlaps implemented in [3]. This may appear counterintuitive at first glance, because the FM index based overlaps particularly cater to short reads, while Minimap2 does not. However, despite holding little promise at first glance, the Minimap2 driven strategy yields drastic improvements in terms of computational expenses, without incurring losses in terms of the quality of the OGs on which strain aware metagenome assembly is based.

Seen from a larger perspective, combining the ideas raised in prior work [3, 4] makes perfect sense; apparently, this insight had passed unnoticed so far. Certainly, a major reason is that [3] had been explicitly designed for fixed ploidy settings, which is rather the opposite of what is needed in metagenome assembly, where numbers and abundances of species and strains are unknown at the beginning. The crucial insight is to realize that the strategy suggested in [3] also works for metagenomes, if presented with well arranged, pre-processed portions of the raw read data.

The last step of StrainXpress (“Global Assembly” in Figure 1) collects all haplotigs into a “master” overlap graph. Construction of such a master overlap graph corresponds to a straightforward operation, since the nodes of the master overlap graph, thanks to the design of the workflow, reflect both strain aware and error corrected sequence.

### Algorithmic Steps: Details

#### Single linkage clustering

For efficiently clustering reads, StrainXpress employs a single linkage clustering algorithm that adopts the successful prior ideas [4]. The overlap file, as generated by Mimimap2, reflects the overlap graph and stores the distance information between each pair of reads. In single linkage clustering, the distance between two clusters is the shortest distance between any member of the first cluster and any member of the second cluster. To quickly determine the corresponding pair of reads in our scenario, StrainXpress sorts the overlap file by the distance scores, such that the “closest” pair of reads (i.e. the pair of reads with the most compatible overlap) is listed at the top, whereas the “most distant” pair of reads (i.e. the pair of reads with the least compatible overlap), is listed at the bottom.

After sorting the overlap file this way, merging clusters in an order that reflects overlap compatibility corresponds to processing the overlap file in one pass, from top to the bottom. The complexity of the corresponding clustering algorithm, including the sorting, amounts to

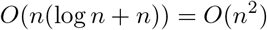

#### Local assembly

The result of single linkage clustering are small groups. It is reasonable to assume that the reads in each of them refer to the one, or at most very little (because likely related) species. StrainXpress assembles the reads in each such group by adopting a strategy suggested by earlier overlap graph based work [3]. This is possible, because numbers of reads per group are sufficiently small.

For substantially increasing speed, StrainXpress makes use of Minimap2 for re-computing overlaps, instead of the FM-index based procedure suggested earlier [3], as already pointed out above.

#### Global assembly

During local assembly, contigs have been generated for each cluster. To further extend these contigs, and identify potential connections across the boundaries of clusters, cluster-specific contigs were collected into a global overlap graph, where vertices correspond to (cluster-specific) contigs and edges correspond to overlap of sufficient quality: an edge corresponds to an overlap of more than 100 bp and identity of at least 0.99, where choices are inspired by analyses presented in earlier work [2, 3].

Corresponding to standard procedures, we further removed all transitive edges, and immediately joined “branch-less” contigs. Here, for identifying branches, we evaluated reads that matched the corresponding contigs. After removal of branches, StrainXpress updates the graph and extends contigs further. This process is repeated iteratively, until no further branches are observed. The final result of this iterative procedure are the “master contigs”, which establish the final output of StrainXpress, ready for usage in downstream analyses.

As pointed out above the Global Assembly step is essentially overlap graph based. However, to further ensure that generation of chimeras is avoided, we only merge contigs into longer, “global” contigs, if the path in the overlap graph leading through these contigs is unique. If the path has branches, hence is not unique, we stop expanding contigs. Note finally that contigs can already be assumed to be strain specific thanks to the procedures that give rise to the earlier steps. Therefore, disturbing effects such as the generation of chimeras remaina minor issue in the Global Assembly step.

#### Sequencing reads: quality control

Before usage, sequencing reads were quality-controlled by *fastp* (version 0.20.1) [6], as a multifunctional FASTQ data preprocessing toolkit. Major functions *fastp* include quality control, detection of adapters, base correction and read filtering.

Bases in the 5’ or 3’ ends of the raw reads exhibiting a Phred score of less than 20, as well as adapters were trimmed. After clipping, only reads longer than 70 bp were kept. Additionally, in the self-overlapping parts of paired-end reads, *fastp* corrects mismatched bases when a high-quality base is paired with a low-quality base.

#### Computation of suffix-prefix read overlaps

StrainXpress utilizes Minimap2 [19] (version 2.18-r1015) for identifying the suffix-prefix overlaps between two reads, as required. In identifying such overlaps, Minimap2 is 3-4 times faster than alternative approaches. While in the earlier approach [4] quality of overlaps was based on a regression scheme that takes error profiles, length and identity of overlap regions into account, StrainXpress calculates an overlap score based on the CIGAR string that is output by Minimap2, see 1 and further explanations below. This eliminates the need for assessing the quality of the overlap across its full length, so comes with considerable gains in terms of runtime.

##### Overlap score

Let *i* be the sequence identity of the overlap region, *o* be the length of the overlap, and *r*_1_, *r*_2_ be the length of the two overlapping reads. We compute the *overlap score D* for the corresponding overlap as

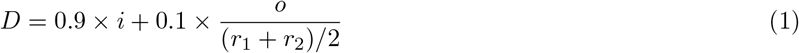

For the sake of illustration, consider that although two reads belong to the same haplotype, the length of their overlap may be short. This requires to strike a balance between length and quality of the overlap. Based on empirical tests, we determined 0.9 (and 0.1 = 1 − 0.9, correspondingly) to work well. While we did not invest in a systematic optimization procedure to determine these parameters, experiments of ours demonstrate that changing these parameters in reasonable ways hardly induces any changes, see Supplementary Table S1 (“Different cluster parameters”). Further, StrainXpress neglects overlaps of length less than 30 bp and identify less than 0.9. These choices are inspired by choices made in earlier work [2, 3].

### Data Sets

#### Simulated Data

To evaluate the performance of StrainXpress, we generated nine synthetic Illumina sequencing data sets using CAMISIM [11] (version 0.0.6), as a popular metagenome simulator. These data sets can be classified into three categories: low complexity data, comprising genomes of 20 strains belonging to 10 species, medium complexity data (100 strains / 30 species) and high complexity data (1057 strains / 376 species). To assess the impact of read length, we generated three data sets for each of the different levels of complexity: 2×250 bp, 2×150 bp, and 2×100 bp, which yielded 9 data sets overall. While the genomes for the “low complexity” and the “medium complexity” data sets were obtained from earlier work [29, DESMAN], the genomes of the high complexity data were downloaded from the 2nd CAMI Human Microbiome Project. See Supplementary Table S1, “Genomes of simulation datasets”, for full information in terms of species and strain content of the data sets referring to the three different levels of complexity. The average coverage per strain in the low complexity, medium complexity and high complexity data sets was 20X, 20X and 10X. As per the principles of CAMISIM, the abundance of different strains is uneven, as sampled from a log-normal distribution.

Further, to evaluate the influence of coverage, we generated 8 data sets that mix simulated and real data. The resulting type of sequencing data set is commonly referred to as “spike-in” data. The idea is to evaluate how methods assemble the simulated, “spiked-in” data—for which one knows the ground truth—as part of a real data scenario. In more detail, we spiked 8 different, real gut metagenome sequencing data sets, resulting from experiments referring to the evaluation of bacteria in stool microbiota relating to the development of eczema [24] (project number: PRJNA272371) with simulated reads from 10 well-known *Salmonella* strains, as downloaded from [27]. For simulating reads from the *Salmonella* strains, we again made use of the CAMISIM simulator. To account for the influence of read coverage, the coverage of the spiked-in strains ranged from 5X to 40X, at steps of 5X, across the 8 real data sets. That is, each of the 8 spiked-in real data sets refers to one particular level of coverage. For details in terms of Genome ID’s and SRA identifiers referring to these data sets, please see Supplementary Table S1, “spike-in salmonella”.

#### Selection of real data sets

We cosidered three real data scenarios in our benchmark experiments:

*Bmock12*, the first sample, reflects a mock community, which includes 12 bacterial strains from 10 species [31]. The read length of *Bmock12* is 2×150 bp (average insert size 302.7 bp), which can be downloaded from the SRA (SRX4901583). The corresponding paired-end reads of 2×150 bp were sequenced using Illumina. Note that the average coverage of *Micromonospora coxensis* is only 0.1X (Supplementary table S1), which hampers any reasonable attempt to assemble that strain. So, one virtually deals with only 11 bacterial strains. The average coverage of the corresponding 11 strains ranges from 74.56X to 3093.79X (median is 1376.35X). For the sake of a less runtime-intense evaluation in the light of the large amount of duplicates among the reads, we randomly extracted 20% of the reads, and further processed only these. Finally, note that challenges of this data set are to assemble the genomes of the two *Marinobacter* and the two *Halomonas* strains, which come at 85% and 99% ANI, respectively.

*NWCs*, the second real data set is a metagenome sequencing data set drawn from natural whey starter cultures [33]. The metagenome samples of NWCs were sequenced using Illumina MiSeq at a read length of 2×300 bp, and, in addition, using PacBio and ONT. As this study focuses on short reads, we did not consider the latter two kinds of reads. In an earlier study, complete genomes for 6 bacterial strains from 3 species were obtained, by means of running a hybrid assembly method [33]. Genbank numbers of the corresponding 6 genomes are CP029252.1, CP031021.1, CP031024.1, CP031025.1, CP029252.1 and CP031021.1; we used these assembled genomes as ground truth when evaluating assembly results. Note that the data set presented itself as unusual, because the reads were pre-trimmed before they were deposited at the SRA. This implies that many reads are of length less than 150 bases, that is much shorter than common Illumina reads. Note further that reads that are too short do not support the identification of patterns of co-occurring variants, which characterize the strains. So, these pre-trimmed reads do not serve the purpose of generating strain aware assemblies—note that including them during assembly led to no improvements neither for StrainXpress nor for the alternative methods, see Supplementary Table S6 and S7; we therefore removed all reads of length less than 150 bases, thereby re-establishing a spectrum of read length that is common for Illumina sequencing experiments. Based on these findings, we generally recommend to remove too short reads before running StrainXpress.

*Gut Metagenome* refers to data stemming from 22 real gut metagenome sequencing data sets, where fragments were sequenced using Illumina HiSeq X Ten, at a read length of 2×150 bp. The foundation of the data sets were 22 samples, each of which refers to one patient either before or after surgery involving thoracic aortic dissection. Each of the 22 patients had gastrointestinal complications [40]. As there was no ground truth readily available, we made use of StrainEst [1] (version 1.2.4), as a tool that depends on the availability of reference genomes to determine genomes at strain level identity from metagenomes (so cannot compete with the tools here, but can be used to generate ground truth for strains that refer to reference genomes of sufficiently high quality). Applying StrainEst to each of the 22 patient samples individually yielded 1899 E. coli strains overall. We then kept only the samples of patients where the raw reads covered more than one of these 1899 strains at at least 95% of their length, which applied for 5 out of the 22 samples. We then further filtered the 1899 *E. coli* strains for those that were sufficiently covered (*>*95% genome length) in any of the 5 patient samples we kept. This yielded 11 strains for which an applicable ground truth was available. For the 5 applicable patient samples, one referred to three, and four referred to two strains. Across the 5 data sets, strains vary in terms of average nucleotide identity (ranging from 96.59 as the least challenging to 98.98 as the most challenging), and depth of coverage per strain ranges from 341X to 17X, see Figure 4 and Supplementary Table S1 (“Gut metagenome”). The SRA identifier of the data set is PRJNA379884; GenBank numbers of the 1899 *E. coli* genomes and SRA number of the 5 assembled samples are listed in Supplementary Table S1. In summary, the 5 data sets just described are meant to reflect a real data based scenario that supports to clearly evaluate how methods behave when varying coverage of strains and divergence between strains.

### Alternative Approaches

We repeat that StrainXpress is novel insofar as methods that decidedly invested in distinguishing strains when assembling metagenomes from short reads had not been available earlier. To nevertheless provide a comparison that appropriately highlights the current status, we considered the following 4 leading state of the art metagenome assemblers: IDBA-UD [28] (version 1.1.3-1), SPAdes [5] (version 3.14.1), MetaSpades, GATB-Minia [8] (version 1.4.1) and MEGAHIT [18] (version 1.2.9). We included SPAdes in addition to MetaSPAdes, because it was recently shown to have advantages over MetaSPAdes with respect to assembling genomes in a haplotype aware manner [2, 4].

Importantly, we recall that all alternative de novo assembly approaches are based on DBGs as an underlying data structure that supports assembly. While IDBA-UD addresses to assemble genomes of uneven coverage (which includes metagenome and single cell sequencing data), SPAdes is a versatile assembler that has shown to prevail also in settings for which it had not been originally designed [2], see also the comment above. Minia is a memory-efficient genome assembler, making use of Bloom filter techniques to represent DBGs efficiently, and serving as the foundation for GATB-Minia, reflecting a pipeline that addresses to assemble metagenomes. MEGAHIT uses concise DBGs to efficiently assemble complex metagenome data sets.

Beyond only considering de novo assembly methods, we also included reference-guided methods in our benchmark experiments, in particular because these reference-guided methods have been designed to assemble genomes in a haplotype-aware manner. As suggested by [12], we considered ConStrains [22], StrainFinder [32], Gretel [26] and DESMAN [29] as currently leading tools. Gretel [26] employs a Bayesian model for local haplotype reconstruction, which uses pairs of single nucleotide polymorphisms (SNPs) showing across several sequencing reads as evidence for haplotypes. DESMAN [29] first assembles reads using MEGAHIT and then proceeds with detecting SNPs in 36 single-copy core genes (“SCGs”) to predict the number and relative abundance of strains. StrainFinder is based on evaluating SNP’s in marker genes (single-copy phylogenetic markers in particular, see [39]) to predict the number of strains and their relative abundance. Similarly, ConStrains is designed to evaluate SNPs in marker genes to predict the number and relative abundance of strains.

### Evaluation Criteria

For comparing methods, we consider genome fraction, NGA50, N50, Error Rate, misassembled contig rate and N rate, as computed by Quast [14] (version 5.0.2). Based on general recommendations, we added the flags –ambiguity-usage all and –ambiguity-score 0.9999 when evaluating metagenomic assemblies. All other parameters reflect default values.

Here, to accurately monitor the performance of the individual methods with respect to strain identity, the reference genome consists of the concatenation of all strain-specific genomes involved in the respective data set one analyzes. In particular, this means that ‘Genome Fraction’, as the percentage of bases in the reference genome against which contigs become aligned, reflects how much of each of the strain-specific genomes can be reconstructed from the reads. In other words, here ‘Genome Fraction’ is the central quantity when assessing strain awareness.

As usual, N50 refers to the length of the shortest contig such that at least 50% of the assembly consists of contigs of that length or greater. NGA50 is the length of the shortest contig such that at least 50% of the true genome (here: the concatenation of all strain specific genomes) are covered by contigs of that length or greater. While the Error Rate is defined as the sum of the mismatch and the indel rate, the N rate is the percentage of ambiguous bases. A misassembled contig is defined to give rise to one of the following scenarios: left and right flanking sequences of the contig (1) both align to the true genome, but leave a gap or overlap themselves by more than 1kbp; (2) align to different strands, or (3) even align to different strains. Identity is calculated by computing the percentage of completely matching bases in the optimal alignment of a contig with the ground truth; note that in general contigs may have several alignments that make sense, while here only the optimal one of them is considered.

## Results

We analyzed the performance of StrainXpress, by comparing it with the available state of the art tools on all of the data sets, all of which we described in the Methods section.

### Experiments

According to the origin of the data, the comparison experiments can be divided into two parts.

#### The first part refers to simulated Illumina data, which includes the spiked-in data sets

Note that the choice of sequencing technology reflects the standards of contemporary short read sequencing based metagenomics. As described above, depending on numbers of species and strains included, we distinguish between data sets of “low complexity”, “medium complexity” and “high complexity”. We also consider the above described “strain-mixing spike-in” data sets, which result from spiking real data with simulated reads generated from known Salmonella strains.

#### The second part refers to the real data sets

As for reference genomes required for evaluating the experiments, we make use of reference genomes when available; as outlined above, reference genomes for ‘Gut Metagenome’ were missing; we recall that we used StrainEst [1] for computing strain specific genomes for five real gut microbiome sequencing data sets that one could use as a ground truth. We also recall that StrainEst depends on the availability of reference genomes at the species level. That implies that StrainEst cannot reasonably compete with the selected methods; however, it is suitable to determine sufficiently reliable ground truth for strains of sufficiently well-studied species.

### Experiments: Simulated Data Sets

We first evaluated all methods, including StrainXpress as well as the 5 alternative state of the art de novo assembly approaches on the synthetic data sets. Please see Table 1 for corresponding results of read length 2×250 bp; see further Supplementary Table 1, “Different length of reads” for results to read lengths of 2×150 and 2×100 bp. See also Supplementary Table 1, “spike-in salmonella” for results on the spike-in *Salmonella* data sets. Quantities referring to the criteria listed in the corresponding Tables were determined using Quast [14], see Methods. We recall that ‘Genome Fraction’, as the percentage of bases of all *strain specific* genomes covered by the assemblies is the immediate, is the central quantity to assess the strain awareness of tools. This explains why we put particular focus on ‘Genome Fraction’ in the description of all experiments that follow.

**Table 1.**
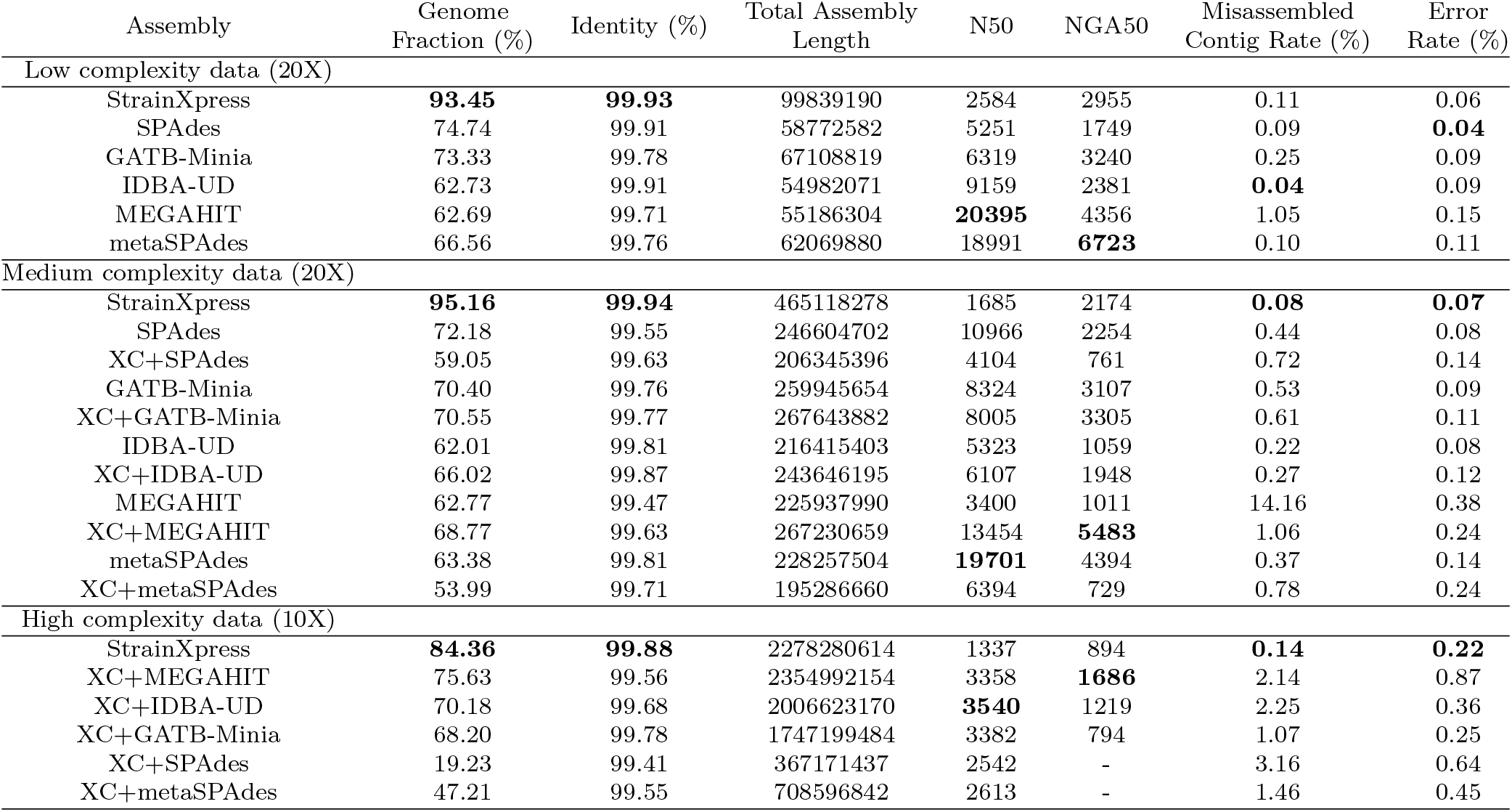
Evaluation of the simulated data (read length 2×250bp). Low complexity data is 20 strains from 10 species, at average coverage 20X per strain. Medium complexity data is 100 strains from 30 species, at average coverage of 20X per strain. High complexity data is 1057 strains from 376 species, at average coverage of 10X per strain. XC is short for the species aware clustering that StrainXpress implements. Identity is the percentage of the completely matching bases of the contig in the optimal alignment with the ground truth. Error Rate = mismatch rate + indel rate.

| Assembly | Genome Fraction (%) | Identity (%) | Total Assembly Length | N50 | NGA50 | Misassembled Contig Rate (%) | Error Rate (%) |
| --- | --- | --- | --- | --- | --- | --- | --- |
| Low complexity data (20X) |  |  |  |  |  |  |  |
| StrainXpress | **93.45** | **99.93** | 99839190 | 2584 | 2955 | 0.11 | 0.06 |
| SPAdes | 74.74 | 99.91 | 58772582 | 5251 | 1749 | 0.09 | **0.04** |
| GATB-Minia | 73.33 | 99.78 | 67108819 | 6319 | 3240 | 0.25 | 0.09 |
| IDBA-UD | 62.73 | 99.91 | 54982071 | 9159 | 2381 | **0.04** | 0.09 |
| MEGAHIT | 62.69 | 99.71 | 55186304 | **20395** | 4356 | 1.05 | 0.15 |
| metaSPAdes | 66.56 | 99.76 | 62069880 | 18991 | **6723** | 0.10 | 0.11 |
| Medium complexity data (20X) |  |  |  |  |  |  |  |
| StrainXpress | **95.16** | **99.94** | 465118278 | 1685 | 2174 | **0.08** | **0.07** |
| SPAdes | 72.18 | 99.55 | 246604702 | 10966 | 2254 | 0.44 | 0.08 |
| XC+SPAdes | 59.05 | 99.63 | 206345396 | 4104 | 761 | 0.72 | 0.14 |
| GATB-Minia | 70.40 | 99.76 | 259945654 | 8324 | 3107 | 0.53 | 0.09 |
| XC+GATB-Minia | 70.55 | 99.77 | 267643882 | 8005 | 3305 | 0.61 | 0.11 |
| IDBA-UD | 62.01 | 99.81 | 216415403 | 5323 | 1059 | 0.22 | 0.08 |
| XC+IDBA-UD | 66.02 | 99.87 | 243646195 | 6107 | 1948 | 0.27 | 0.12 |
| MEGAHIT | 62.77 | 99.47 | 225937990 | 3400 | 1011 | 14.16 | 0.38 |
| XC+MEGAHIT | 68.77 | 99.63 | 267230659 | 13454 | **5483** | 1.06 | 0.24 |
| metaSPAdes | 63.38 | 99.81 | 228257504 | **19701** | 4394 | 0.37 | 0.14 |
| XC+metaSPAdes | 53.99 | 99.71 | 195286660 | 6394 | 729 | 0.78 | 0.24 |
| High complexity data (10X) |  |  |  |  |  |  |  |
| StrainXpress | **84.36** | **99.88** | 2278280614 | 1337 | 894 | **0.14** | **0.22** |
| XC+MEGAHIT | 75.63 | 99.56 | 2354992154 | 3358 | **1686** | 2.14 | 0.87 |
| XC+IDBA-UD | 70.18 | 99.68 | 2006623170 | **3540** | 1219 | 2.25 | 0.36 |
| XC+GATB-Minia | 68.20 | 99.78 | 1747199484 | 3382 | 794 | 1.07 | 0.25 |
| XC+SPAdes | 19.23 | 99.41 | 367171437 | 2542 | - | 3.16 | 0.64 |
| XC+metaSPAdes | 47.21 | 99.55 | 708596842 | 2613 | - | 1.46 | 0.45 |

Because the reference-guided alternative approaches that we considered (Gretel, DESMAN, StrainFinder and ConStrains) required particular treatment in terms of additional, special data sets, they could not compete with the de novo assembly tools in the frame of an overall comparison. We therefore discuss the results referring to the reference-guided tools in an additional, separate paragraph “Comparison with reference-guided methods”, see further below.

#### Low Complexity Data: Speed and Strain Awareness at Coverage of 20X

We recall that the ‘low complexity’ synthetic data set contains 20 strains belonging to 10 species, at an average coverage of 20X per strain.

##### Part I: Computational efficiency of overlap graph based strategy

StrainXpress is based on overlap graphs (OGs), which are known for being computationally intense from a theoretical point of view. Getting OG based approaches to work sufficiently fast in practice is meant to be a major argument of this study. We therefore first demonstrate that StrainXpress indeed is sufficiently fast when being used in realistic scenarios. For that, note further that the complexity of the data, the number of strains, as well as their identity and coverage do not have much of an influence on the computational bottlenecks that OG based approaches can struggle with. Therefore, in order to run a resource friendly analysis, we solely focus on the low complexity data set (2×250 bp) when assessing the speed. In fact, we observed similar requirements also for the other data sets (without that we provide full details).

The first part of our analysis of runtime and memoryy requirements refers to comparing StrainXpress with the original OG based strategies that provided inspiration for StrainXpress. Because the original strategies were known to be very slow, StrainXpress is meant to establish an improvement over these original strategies. Here, we show that StrainXpress is faster by (a whopping) 1-2 orders of magnitude. We further dissect consumption of runtime and memory into the parts that refer to the different stages of StrainXpress, ‘Clustering Reads’, ‘Local Assembly’ and ‘Global Assembly’, see Figure 1.

Please see Supplementary Tables S2 and S3 for the following results. One observes that ‘StrainXpress - Clustering’ is 46 times faster than the earlier work [4] (Supplementary Table S2) without any noticeable losses in terms of assembly quality (compare ‘StrainXpress’ and ‘OGRE + Local Assembly’ in Supplementary Table S3). Importantly, StrainXpress yields 899 clusters, instead of only 568 clusters, which further enhances speed because of enhanced parallelization.

Also, ‘Local Assembly’ is 8 times faster than the original template [3] (see Supplementary Table S2), again without remarkable losses in terms of assembly quality: note that although Genome Fraction drops by one point (Supplementary Table S3), Genome Fraction still exceeds all alternative methods by nearly 20%, see the discussion below and Table 1.

Finally, note that runtime advantages persist when replacing the components of StrainXpress with the precursor strategies, both one by one, and in combination.

##### Part II: Recovery of 20% additional strain specific sequence

See Table 1 (read length 2×250 bp) for the following. The most relevant observation, because immediately referring to strain awareness, is that StrainXpress raises Genome Fraction by at least 20% in comparison with the other methods (StrainXpress: 93.45% versus 74.74% by SPAdes, as the second highest scoring). Arguably, this supports the claim that StrainXpress is the only metagenome assembler that is sufficiently strain aware.

Further, StrainXpress ranks first in terms of identity (StrainXpress:99.93% vs. IDBA-UD: 99.91%), second for Error Rate (StrainXpress: 0.06% vs. SPAdes: 0.04%) and fourth for Misassembled Contigs Rate (StrainXpress: 0.11% vs. IDBA-UD: 0.04%); all of these reflect very low numbers in general. The N50 being smaller than for other methods is put into perspective by the NGA50 which unlike N50 refers to the reference sequences being roughly on a par for all methods, including StrainXpress.

While the contigs of StrainXpress are shorter relative to the length of the assembly (as measured by N50), they are not when comparing the length of the contigs with respect to the true genomes that needed to be reconstructed. On that point, note that the length of the assembly of StrainXpress is substantially longer than the ones delivered by the alternative methods, which puts the small N50 into perspective. On a side remark, MEGAHIT outperforms all other methods in terms of contig length (N50 in particular), obviously traded for a larger amount of misassemblies (exceeding the rate of others by one order of magnitude).

Please see the Supplementary Table S1, Genome Fraction is reduced for all methods except MEGAHIT as read length decreases to 2×150 or 2×100 bp. However, StrainXpress still outperforms other methods. At read length of 2×150 bp, StrainXpress (Genome Fraction: 89.14%) reconstructs 25.47% more of the strain-specific genomes than the second ranked (Genome Fraction: 63.67%). Decreasing the read length to 2×100 StrainXpress comes out at 76.51% Genome Fraction in comparison to 63.23% Genome Fraction achieved by the second ranked MEGAHIT, so raises Genome Fraction by at least 13.28% in comparison with other methods. StrainXpress further outperforms other assemblers in terms of identity, Error Rate and Misassembled Contigs Rate.

#### Medium complexity data: full strain awareness, no errors and no misassemblies

See Table 1 (read length 2×250 bp) for the following. We recall that the medium complexity data set contains 100 strains from 30 species, at average coverage 20X per strain. We considered the application of StrainXpress’s clustering procedure (referred to as ‘XC’ in Table 1) as a pre-processing step for the other methods (including SPAdes). We recall that prior, taxonomy aware clustering of raw metagenome data was suggested as an interesting general strategy earlier [4]. Remarkably, this led to considerable improvements for IBDA-UD and MEGAHIT, while affecting SPAdes to the worse (and hardly affecting GATB-Minia). We refer to the Supplement for a detailed evaluation with respect to how using XC affects the qualities of the assemblies of alternative methods.

StrainXpress achieves nearly full strain awareness also on this data set of greater complexity, improving over other methods by large margins (95.16% vs. 72.18% by SPAdes). Interestingly, StrainXpress assemblies now also contain the least errors (0.07% vs. 0.08% by SPAdes) and StrainXpress seems to be the only method that can preserve the low misassembly rate (0.08% vs. 0.22% by IBDA-UD) in the light of this more complex data, with all other methods experiencing at least 2 times more misassemblies. Again, StrainXpress having small N50 is put into perspective with the NGA50 not being inferior with respect to others, and compensated by the much reduced amount of misassemblies. In summary, also on data sets of such characteristics, StrainXpress appears as the only tool that can hold the promise of delivering strain aware assemblies.

At read length of 2×150 or 2×100 bp, StrainXpress still outperforms the other methods in terms of all Genome Fraction, identity and misassembled contig rate, see Supplementary Table S1. At read length of 2×150 bp, StrainXpress (Genome Fraction: 81%) reconstruct 18.93% more of the true genomes than the second ranked MEGAHIT (Genome Fraction: 62.07%). When decreasing the read length further to 2×100 bp, the Genome Fraction achieved by StrainXpress is reduced to 67.67%. Still, this outperforms the second ranked method by 7.59% (MEGAHIT: 60.08%).

#### High complexity data: reliable reconstruction of strains of coverage as low as 5X

We recall that the data set reflects 1057 strains from 376 species, with strains sequenced at an average coverage of 10X. For read length 2×250 bp, for example, this amounted to 62 131 250 paired-end reads overall (and, naturally, even more for read lengths of 2×150 bp and 2×100 bp). Due to the huge amount of reads, StrainXpress is the only tool that is able to process this data set without prior treatment. This highlights the practical usefulness of the divide-and-conquer strategy, and provides further evidence for the computational efficiency of StrainXpress in practice.

To nevertheless make it possible to run the other tools, and provide a meaningful comparison, we ran the alternative approaches on the clusters that were generated during the first stage (which we continue to refer to as ‘XC’). In that, we profited from the insight having gained from the experiments on the medium complexity data. Note, however, that for medium complexity prior treatment was an option, while here, for high complexity, prior treatment is a necessity. We remind that on the medium complexity data prior treatment led to improvements for all alternative approaches apart from SPAdes.

See Table 1 for results on read length 2×250 bp. StrainXpress outperforms all other methods by large margins in terms of Genome Fraction, Error and Misassembly Rates. As usual, StrainXpress has smaller N50, compensated by NGA50 that is on par with those of other tools.

In comparison with earlier results, StrainXpress does no longer achieve Genome Fraction of more than 90%. See Figure 2 for the corresponding explanation: for strains of coverage less than 5X, StrainXpress achieves Genome Fraction of a bit more than 60%. For coverage of 5X to 10X, StrainXpress again successfully assembles nearly 90% of the strains. Genome Fraction rises to more than 95% on average for strains of coverage of least 10X. Across all coverage ranges, StrainXpress outperforms all other tools significantly. For 5X and more, StrainXpress is also significantly more stable, as indicated by smaller sized boxes in Figure 2. This last point suggests that strain specific sequential phenomena have less of an influence on the performance of StrainXpress than that of the other tools.

**Figure 2.**
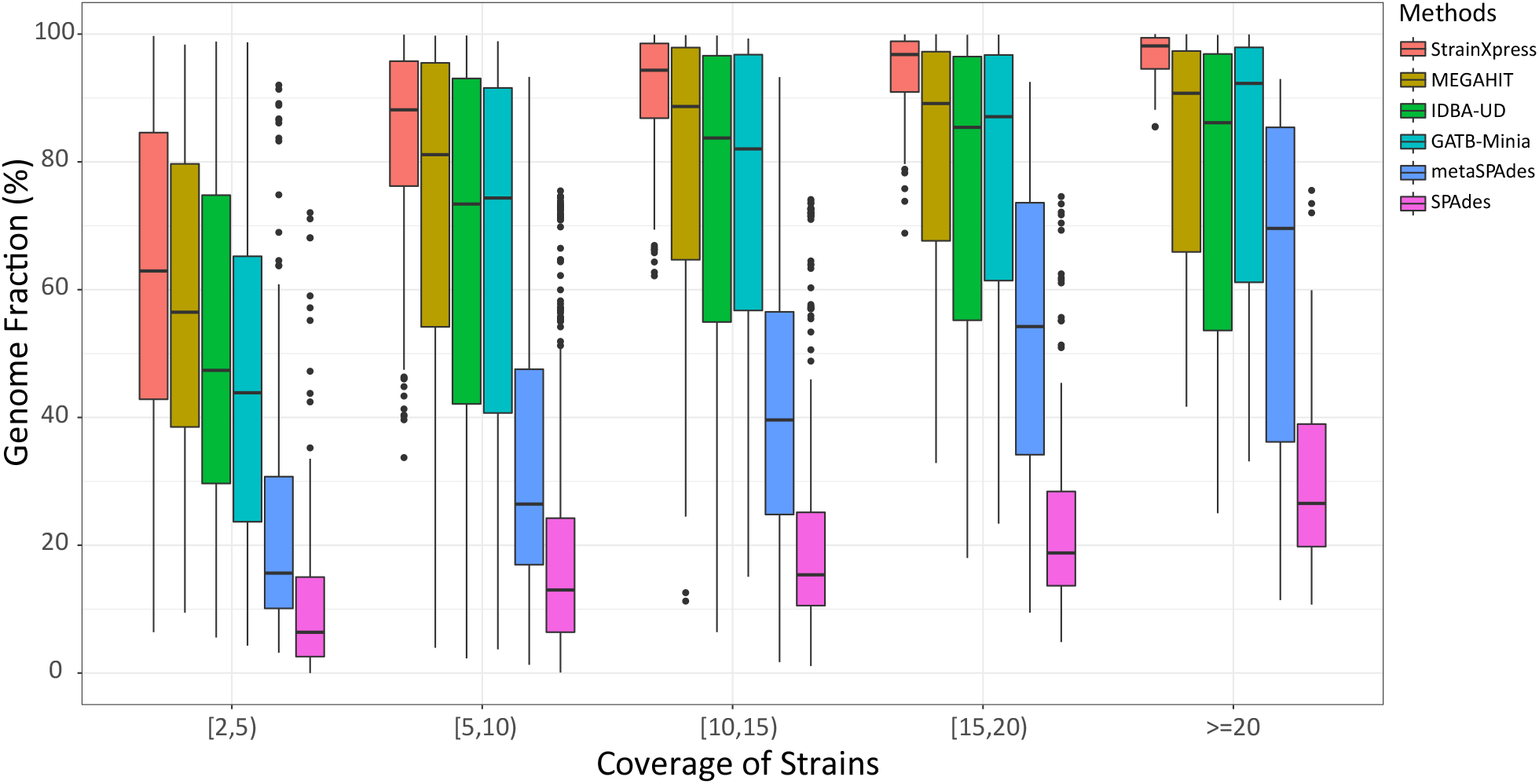
Genome fraction versus the coverage of strains on the high complexity data set (2×250bp). The high complexity data set contains 1057 strains from 376 species. The average coverage of the 1057 strains is 10X, but varies according to a log-normal distribution. We display the genome fraction of the different strains in the different coverage intervals. Different colors denote different assembly methods.

For results relating to shorter read lengths (2×150 bp, 2×100 bp), please see Supplementary Table S1. For 2×150 bp, StrainXpress still outperforms the other methods in terms of the central measures Genome Fraction, Identity and Misassembled Contig Rate. For read length of 2×100 bp, however, Genome Fraction of XC+IDBA-UD (52.77%) is slightly higher than that of StrainXpress (50.38%). To analyze this further, we found that decreasing the length of the contigs that are evaluated by Quast (which operates at a default of 500 bp) to 400 bp or 300 bp—which is still considerably longer than the original read length—we find that Genome Fraction increases for StrainXpress (400 bp: 56.34%, 300 bp: 63.34%), while not noticeably increasing for the other tools (XC+IBDA-UD – 400 bp: 53.77%, 300 bp: 55.04%). Similar effects show for reads of length 2×150 bp already. This indicates that StrainXpress still generates substantial amounts of high-quality contigs, a substantial portion of which is shorter than 500 bp, however. So, comparing tools across the whole spectrum of contig length, StrainXpress still has considerable advantages. The likely reason for this phenomenon is that due to the short read length, branches in the assembly graph show earlier, which prevents expanding contigs further.

Moreover, note that the strain composition for certain species in the high complexity data is extremely challenging. For example, the 71 *Neisseria meningitidis* strains come at ANI between 97.76% and 99.9997%, at a median of 99.97%, which renders them extremely difficult to distinguish. This further emphasizes the difficulties above described, because branches in the assembly graph tend to show earlier for such species.

#### Strain-mixing spike-in data: the influence of coverage

We assembled the 8 “strain-mixing spike-in” data sets and evaluated the quality of the assemblies of the 10 spiked in *salmonella* strains using metaQuast. We recall that an important particular aspect was that the 8 data sets varied in terms of coverage of the spiked-in strains, ranging from 5X to 40X in steps of 5X, across the 8 data sets. See Figure 3 for results with respect to Genome Fraction and see Supplementary Table S1 for full details. As becomes obvious from Figure 3, StrainXpress still reconstructs 70.26% of the genomes at coverage of only 5X, which outperforms the second best method by 22% (MEGAHIT: 48.26%). From coverage of 15X onwards, the Genome Fraction increases to 84.29% and tends to stabilize. In comparison with other methods, this means an advantage of at least 36.05% (second best: MEGAHIT at 48.24%). As becomes further evident from Supplementary Table S1, StrainXpress achieves competitive, if not even optimal performance, also in the categories ‘Misassembled Contigs’, ‘Identity’ and ‘Error Rate’. In a final remark, note that the effects observed generally agree with those observed for the high complexity data set (see Figure 2).

**Figure 3.**
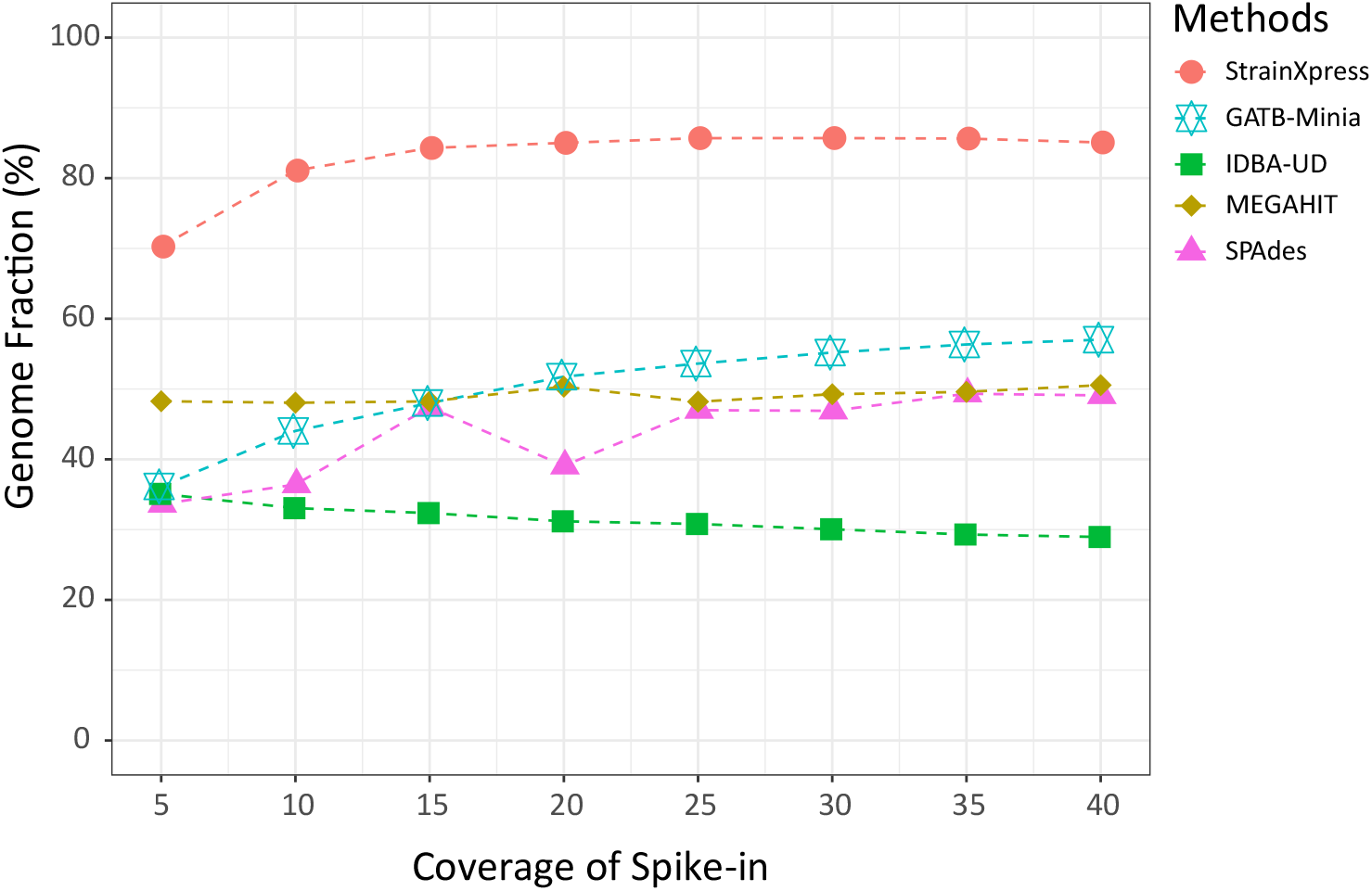
We generated simulation data sets with different coverage, which contains 10 *salmonella* strains. The synthetic reads are mixed with real gut metagenome sequencing data and then assemble them with different approaches. The figure presents the change of Genome Fraction in distinct assembly methods with the increase of coverage of the ten *salmonella* strains.

#### Comparison with reference-guided methods

In the following, we discuss the experiments run for the above-mentioned 4 reference-guided tools. We recall that the 4 methods were suggested by [12] as establishing the current state of the art.

##### Gretel

Initially, we tried to run Gretel on the low complexity data set containing reads of length 2×250 bp. Following the instructions provided on Gretel’s website (https://gretel.readthedocs.io/en/latest/protocol.html), we first processed the low complexity data using SPAdes and MEGAHIT, as the currently most popular state-of-the-art methods for de novo metagenome assembly (recalling that SPAdes outperforms MetaSPAdes, which explains our preferences), and used the resulting contigs as reference for Gretel. However, Gretel did not terminate within two weeks (on 32 CPU’s and 500GB RAM), such that we aborted the run.

To nevertheless provide a meaningful comparison, we generated a smaller data set of even lower complexity. This small data set contains 3 *Salmonella* strains drawn from [27], each of which comes at a coverage of 20X (read length again is 2×250 bp). Please see Supplementary Table S1 for Genome Id’s and ANI’s.

For the results obtained, please see Supplementary Table S8. MEGAHIT+Gretel slightly outperforms StrainXpress in terms of Genome Fraction (StrainXpress: 90.84%; SPAdes+Gretel: 83.78%; MEGAHIT+Gretel: 92.54%). However, this (slight) advantage comes at enormous expenses in terms of Duplication Rate (StrainXpress: 1.21; SPAdes+Gretel: 8.16; MEGAHIT+Gretel: 9.66), Error Rate (StrainXpress: 0.059; SPAdes+Gretel: 1.164;

MEGAHIT+Gretel: 1.861), Identity (StrainXpress: 99.93; SPAdes+Gretel: 97.70; MEGAHIT+Gretel: 97.12) and Misassembled Contig Rate (StrainXpress: 0.06; SPAdes+Gretel: 0.46; MEGAHIT+Gretel: 0.68). Moreover, MEGAHIT+Gretel and SPAdes+Gretel are 45 times slower than StrainXpress.

##### StrainFinder

We did not manage to run StrainFinder nor on our regular benchmark data neither on the small data set generated for Gretel due to an error thrown during the pre-processing step. This agrees with the experiences of other users. See the Supplement (“Alternative Methods: Errors”) for more details.

##### ConStrains

Just as StrainFinder, also ConStrains throws an error on any of the data sets considered. Again, analogous issues were also encountered by other users and, unfortunately, have so far (as of March 20, 2022) not been commented on by the authors. Again, see the Supplement (“Alternative Methods: Errors”) for more details.

##### DESMAN

Unlike StrainFinder and ConStrains, DESMAN successfully terminated on the small data set that we had generated for running Gretel. However, DESMAN solely estimates the number of strains and their relative abundances. It does not provide sufficient haplotype-specific information based on which one can generate non-ambiguous, haplotype-aware contigs. This does not allow one to establish a meaningful comparison in terms of metagenome *assembly* ; evidently, DESMAN is not primarily meant to be an assembly tool. Of course, this does not deny its obvious usefulness when evaluating the strain content of metagenomes in terms of other relevant quantities, apart from reconstructing the genomes of strains.

##### Summary

Unlike the de novo assembly approaches, all reference-guided methods either encountered difficulties when processing both our simulated and real data sets or are not designed to compute assemblies. We feel that it is important to mention that these tools may put other valuable challenges that arise in metagenome analysis, such as estimating numbers and abundances of strains and species, in their main focus.

#### Experiments: Real Data Sets

We further evaluated all approaches on the real data sets ‘Bmock 12’, ‘NWCs’ and ‘Gut metagenome’.

#### Bmock12: Successful separation of strains of 99% identity

We recall that the ‘Bmock12’ data set contains 11 strains from 9 species, so is of rather low complexity. The two species that exhibit more than 1 strain are *Marinobacter* and *Halomonas*. Naturally, we will pay particular attention to them in the following.

See Table 2 for the following results. All methods achieve fairly large Genome Fraction, which is explained by the low complexity of the data: 7 out of 9 species only have 1 strain, so can be assembled without difficulties regardless of the approach. Nevertheless, the Genome Fraction of StrainXpress exceeds that of others by at least 3.7%, approaching 99.04% overall, which demonstrates near-perfect separation of all strains involved.

**Table 2.** Evaluation of the real data sets. Having removed a strain of coverage too low to allow for reconstruction, Bmock12 effectively corresponds to 11 effective bacterial strains. NWCs reflects metagenome sequencing data of natural whey starter cultures (NWCs), and contains 6 bacterial strains from 3 species. Error Rate = mismatch rate + indel rate. For results on Gut Metagenome, please see Figure 3.

| Assembly | Genome Fraction (%) | Identity (%) | Total Assembly Length | N50 | NGA50 | Misassembled Contig Rate (%) | Error Rate (%) |
| --- | --- | --- | --- | --- | --- | --- | --- |
| Bmock12 |  |  |  |  |  |  |  |
| StrainXpress | **99.04** | 99.93 | 55332069 | 60566 | 65743 | 0.78 | 0.018 |
| GATB-Minia | 95.31 | 99.92 | 49058237 | 96537 | 80434 | 0.47 | 0.014 |
| SPAdes | 95.28 | 99.98 | 49055870 | **189251** | **171570** | 0.06 | 0.012 |
| metaSPAdes | 94.55 | 99.97 | 48998826 | 171793 | 155762 | 0.13 | 0.028 |
| IDBA-UD | 94.67 | **99.99** | 48465926 | 72765 | 60987 | **0.05** | **0.006** |
| MEGAHIT | 93.25 | 99.87 | 48637140 | 120129 | 105626 | 2.79 | 0.027 |
| NWCs |  |  |  |  |  |  |  |
| StrainXpress | **75.29** | 99.47 | 8858666 | 1056 | 636 | 3.34 | 0.30 |
| SPAdes | 59.37 | 99.38 | 6083388 | 10160 | - | 2.72 | 0.08 |
| metaSPAdes | 57.96 | **99.68** | 5767394 | 9871 | - | **1.05** | **0.05** |
| MEGAHIT | 57.81 | 97.78 | 6141276 | **14456** | - | 12.52 | 0.16 |
| GATB-Minia | 56.78 | 98.78 | 5779411 | 11081 | - | 4.16 | **0.05** |
| IDBA-UD | 56.44 | 98.87 | 5873327 | 9320 | - | 3.97 | 0.07 |

To evaluate the corresponding details, we further stratified the results by the individual strains; see Table 3. The two *Marinobacter* strains agree on 85% of their nucleotides (average nucleotide identity (ANI) = 85%), whereas the two *Halomonas* strains have ANI = 99%, exposing the *Halomonas* strains as the real challenge. We recall that the 7 strains of the other species pose no difficulties for any method. While StrainXpress has relatively small advantages on *Marinobacter* (StrainXpress: 99.5% vs. other Methods: 97-98%), StrainXpress has decisive advantages over all other approaches on *Halomonas*: StrainXpress reconstructs both strains at at least 95.5%, whereas other methods do not exceed 70% (second best: GATB-Minia) for at least one of the strains. In summary, StrainXpress is the only method that can successfully distinguish between strains of identity of 99%. This demonstrates clear advantages relative to strain specific reconstruction of genomes.

**Table 3.**
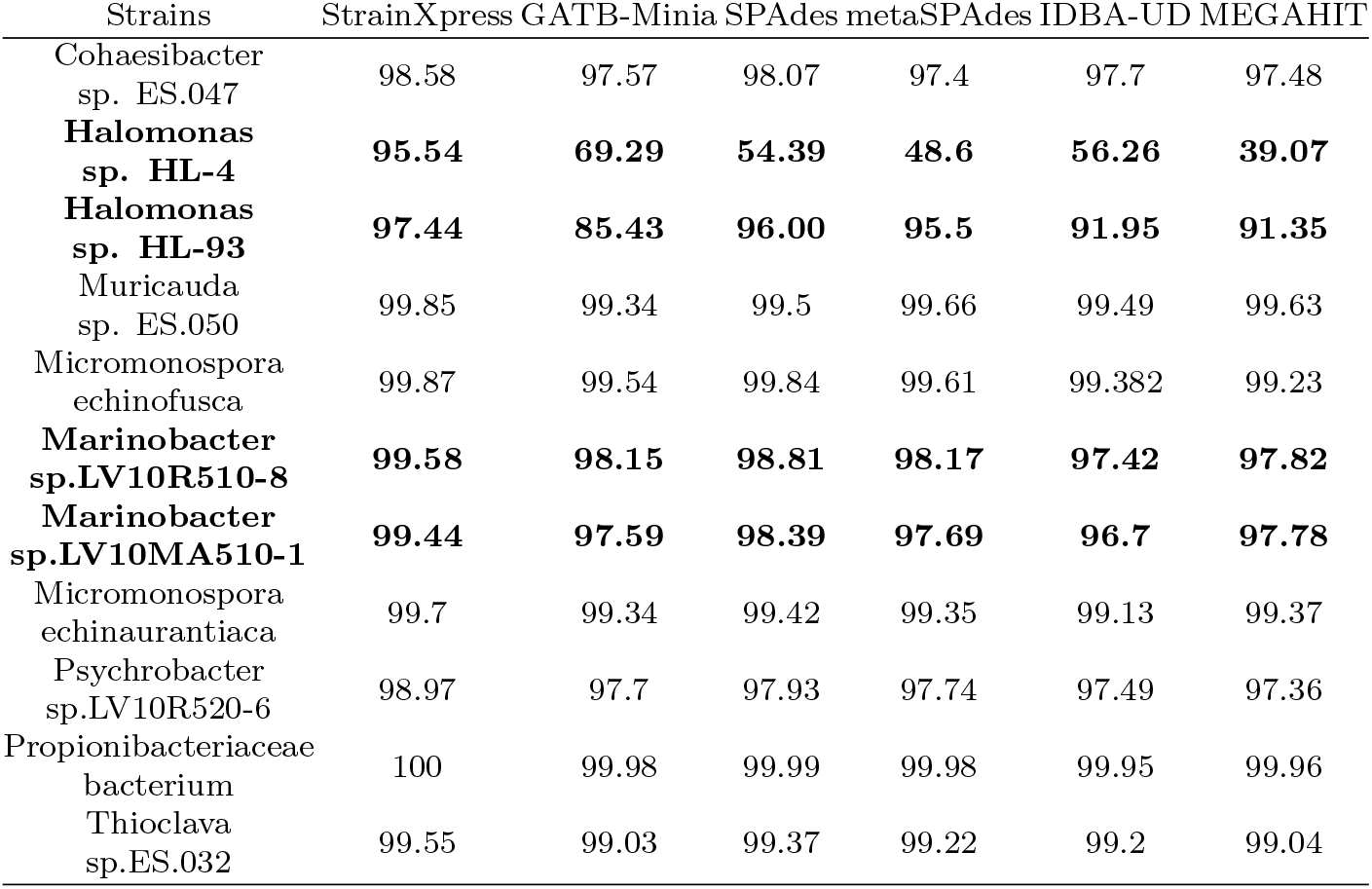
Genome Fraction of individual strains in the Bmock12 data set of the assemblies of the different approaches.

As for N50 and NGA50, SPAdes clearly outperforms all other tools, tripling the values of StrainXpress in particular. Of further note, one sees that in addition to SPAdes (0.06%), also IBDA-UD (0.05%) achieves a remarkably low misassembly rate. Although still very low (that is below 1), StrainXpress has slight disadvantages here (Misassembled Contig Rate: 0.78%). As always, the explanation for this is the fact that StrainXpress does not collapse sequence patches into longer consensus sequence as easily as the other methods.

#### NWCs: Dominant strain awareness on highly similar strains above 25X

This data set contains 3 species (Streptococcus thermophilus, Lactobacillus delbrueckii, Lactobacillus helveticus), of 2 strains each, coming at ANI’s of 99.99%, 99.24% and 98.03%, respectively, rendering this data set particularly challenging in terms of strain diversity.

Note that we removed reads that became too short by the trimming procedure that was originally applied (see Methods for details). It is important to understand that beyond just re-establishing a data set following a common read length distribution, the corresponding reduction in terms of coverage rendered the data set particularly challenging: the average coverage of strains was 25.47X, at a minimum of 7.46X for one of the Lactobacillus helveticus strains. So the reduction in terms of reads allowed to evaluate tools with respect to low sequence coverage when dealing with real data.

Indeed, we found this data set to be the most difficult one for StrainXpress: while still showing ∼15% more Genome Fraction than all competitors (see Supplementary Table S7), Genome Fraction drops to 76.13% in the NWCs data set. For explaining these effects, we stratified the fraction of reconstructed genomes relative to the individual strains, see Supplementary Table S7. We see that Genome Fraction drops from 97.28% and 94.57% for the Streptococcus thermophilus strains (38.36X and 37.42X), to 68.71% and even 39.59% for the Lactobacillus helveticus strains (13.07X and 7.46X). In summary, results point out that StrainXpress requires approximately 30X for strains that are highly similar to other strains, to operate at the usual levels of quality, in this real scenario, while clearly dominating the other approaches on all strains.

#### Gut Metagenome: Distinction of near-identical strains at low coverage

We recall that ‘Gut Metagenome’ consists of 5 data set that refer to 11 *E. coli* strains overall (see Methods for full details). While the average of coverage of the 11 strains is 76X (17X ∽341X), the average ANI between two strains stemming from one of the 5 samples is 97.97% (96.59 - 99.65%). Reads were assembled for each of the 5 data sets individually. According to the variations in terms of coverage and ANI of strain content, the data sets correspond to 5 real scenarios of varying degrees of difficulty, so enable the evaluation of the limits of the methods in terms of strain specific reconstruction of genomes relative to identity of strains and coverage.

In the following, we focus on evaluating Genome Fraction, as the decisive quantity in the context of these data sets; see Figure 4 for corresponding results, where strains are listed in descending order relative to the depth of coverage at which they were sequenced. Note however that, in addition to coverage, average nucleotide identity (ANI) is an important factor, too. One observes that Genome Fraction increases on increasing coverage per strain, which was to be expected. StrainXpress achieves 73.41% for the strain of lowest coverage (17X) while achieving 98.04% for the strain of greatest coverage (341X). We recall that every strain has at least one other strain that matches it closely, pointing out that the performance rates of StrainXpress are very competitive. Indeed, all other methods only reach 10-25% Genome Fraction for at least one of the lowly covered strains. Also, StrainXpress’ Genome Fraction drops below 75% only for strains of coverage below 24X, whereas all other methods require at least 50X for successful reconstruction of strain-specific genomes.

**Figure 4.**
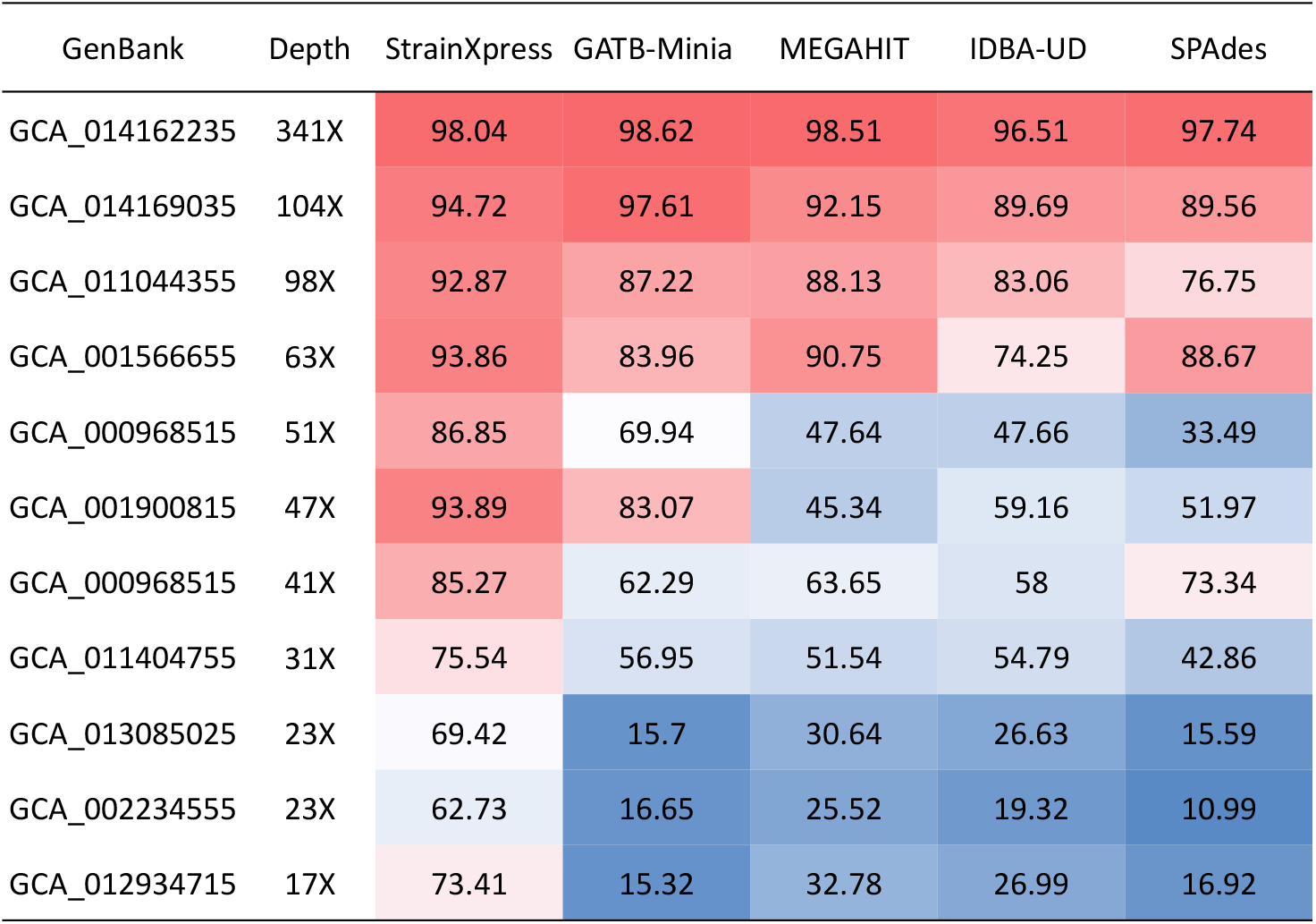
Genome Fraction (%) of “Gut Metagenome”. In the 5 real gut metagenome sequencing data, StrainEst predicted 11 strains applicable to serve as ground truth. The performance of the different methods is evaluated with METAQUAST. The first column displays the GenBank access numbers of the 11 strains. Numbers in the heatmap correspond to genome fraction.

See Supplementary Table S1 for a full overview of results, stratified by the different data sets to which the strains belong. As for criteria other than Genome Fraction, one can see in Supplementary Table S1 that the usual trends continue also here: StrainXpress’ contigs are shorter in comparison with other approaches. This is compensated, however, by less misassemblies: obviously, StrainXpress does not stitch together sequence patches from different strains, as a potential explanation for the superior Misassembled Contig Rate. Also, while exhibiting excellent Error Rate, StrainXpress does not necessarily dominate the other approaches (in particular SPAdes) in terms of this particular category.

In summary, StrainXpress clearly outperforms all other methods when reconstructing strains across all combinations of coverage and identity of strains reflected by the data sets presented.

## Discussion

Although different strains from the same species can be near-identical in terms of sequence, they can vary greatly in terms of their phenotypes, such as drug resistance or pathogenicity. Short read sequencers, which in the meantime belong to the standard equipment in genomics laboratories, offer inexpensive means to analyze the sequence content of mixed samples such as metagenomes.

This combination explains why *assembling strain specific genomes from metagenome short read data* is of utmost current interest. On the one hand, it is worthwhile to study the spectrum of phenotypes characterizing a metagenome by way of a routine experimental protocol. On the other hand, however, the technical challenges are enormous: strain aware assembly of metagenomes has remained a largely unresolved problem, despite the well-deserved attention it has been receiving throughout.

Here, we have presented StrainXpress, which implements a major step towards resolving the issue. In benchmarking experiments of great variability that adhere to the currently highest standards, we have demonstrated that StrainXpress outperforms the current state of the art by large margins. StrainXpress demonstrated its greatest advantages when differences between strains were small, or when strains were subject to low read coverage.

Overall, StrainXpress appeared to be the only approach whose assemblies tended to cover at least 90% of strain specific sequence content. Toughest competitors have been trailing by at least roughly 20%, on all data sets. Only when undergoing newly developed preprocessing routines—themselves being integral components of StrainXpress—losses in performance could be limited to below 20% for the prior approaches.

StrainXpress is a *de novo* approach. As such, it does not suffer from reference-induced biases, so has relevant advantages over reference-assisted approaches. Importantly, reliable reference genomes are not yet available for a multitude of prokaryotic species, which prevents the usage of reference genomes altogether. This justifies why *de novo* assembly is regarded the approach of choice when distinguishing strains in metagenomes. Beyond this theoretical insight, usage of *de novo* assembly approaches became further justified through experiments with reference-guided tools. Unlike the de novo approaches, reference-guided tools often tend to focus on challenges other than assembly when analyzing metagenomes in terms of species and strain content.

Key to success for the design of StrainXpress was the insight that recent progress in overlap graph based short read assembly had pointed out a way to successfully identify the genomes of strains, and not just their species. The foundation of the corresponding ideas are the fact that patterns of co-occurring mutations—as the decisive characteristic sequential phenomenon of strains—can be highlighted optimally with overlap graphs. Unlike de Bruijn graph based approaches, they do not chop reads into smaller subsequences which breaks up, and therefore masks such patterns.

While this establishes a clear conceptual advantage, the decisive challenge when dealing with overlap graph based approaches are requirements in terms of computational resources. In fact, the computational “heaviness” required to both design a framework in which the different subroutines could be integrated, and to develop and implement practical solutions through which the framework and its integral components were applicable in real world scenarios.

The combination of framework overall, and sufficiently lightweight integral components that support it, has established a methodical novelty in metagenome assembly. We have demonstrated that our implementation of substantially accelerated individual components has led to very feasible run times in practice, at no sacrifices in terms of the quality of the assemblies. The general flexibility (and, arguably, the general value) of our approach was further highlighted by the fact that one of the individual components can also be used to improve the results of alternative approaches, regardless of the methodical details of the approaches, and to enable them to work with much larger data sets.

In conclusion, we have presented StrainXpress, a *de novo* method that assembles individual genomes from metagenomes at strain resolution. In this, StrainXpress not only is the first fully overlap graph based approach that overcomes the underlying great technical challenges. It is also the first approach to redeem the great advantages that such overlap graph based approaches had recently been promising. In summary, our approach is able to reconstruct substantially greater portions of individual genomes at strain resolution, with advantages being most striking when dealing with extremely similar strains, or strains lacking sequencing coverage.

Further improvements are conceivable. In particular, the short length of Illumina type reads puts constraints on the length of the contigs a strain aware assembler can compute. This prevents to compute strain specific contigs that span the genomes of the individual strains at their full length: trying to increase their length introduces ambiguities that cannot be resolved using short reads. For this issue, third generation sequencing reads used in the context of overlap graphs present promising opportunities. The great Error Rate of third generation sequencing reads, however, introduce a range of issues that still need to be resolved.

For the time being, StrainXpress, the approach we have presented, points out a way to thoroughly explore the strain content of huge amounts of metagenomes that so far have been (and further will be) sequenced using Illumina type platforms.

## 1 AVAILABILITY

The source code of StrainXpress is https://github.com/kangxiongbin/StrainXpress.

## Acknowledgments

We would thank Jasmijn Baaijens and Marleen Balvert for their insightful discussions.

## Footnotes

1 In the sense of sharing a large part of their nucleotides in identical order, which applies for the genomes of two different strains of the same species metagenome analysis.

## References

1. D. Albanese and C. Donati. Strain profiling and epidemiology of bacterial species from metagenomic sequencing. Nature communications, 8(1):1–14, 2017.

2. J. A. Baaijens, A. Z. El Aabidine, E. Rivals, and A. Schönhuth. De novo assembly of viral quasispecies using overlap graphs. Genome research, 27(5):835–848, 2017.

3. J. A. Baaijens and A. Schönhuth. Overlap graph-based generation of haplotigs for diploids and polyploids. Bioinformatics, 35(21):4281–4289, 2019.

4. M. Balvert, X. Luo, E. Hauptfeld, A. Schönhuth, and B. E. Dutilh. Ogre: Overlap graph-based metagenomic read clustering. Bioinformatics, 37(7):905–912, 2021.

5. A. Bankevich, S. Nurk, D. Antipov, A. A. Gurevich, M. Dvorkin, A. S. Kulikov, V. M. Lesin, S. I. Nikolenko, S. Pham, A. D. Prjibelski, et al. Spades: a new genome assembly algorithm and its applications to single-cell sequencing. Journal of computational biology, 19(5):455–477, 2012.

6. S. Chen, Y. Zhou, Y. Chen, and J. Gu. fastp: an ultra-fast all-in-one fastq preprocessor. Bioinformatics, 34(17):i884–i890, 2018.

7. H. Cheng, G. T. Concepcion, X. Feng, H. Zhang, and H. Li. Haplotype-resolved de novo assembly using phased assembly graphs with hifiasm. Nature Methods, 18(2):170–175, 2021.

8. R. Chikhi and G. Rizk. Space-efficient and exact de bruijn graph representation based on a bloom filter. In International Workshop on Algorithms in Bioinformatics, pages 236–248. Springer, 2012.

9. K. Clarke, Y. Yang, R. Marsh, L. Xie, et al. Comparative analysis of de novo transcriptome assembly. Science China Life Sciences, 56(2):156–162, 2013.

10. N. Fierer. Embracing the unknown: disentangling the complexities of the soil microbiome. Nature Reviews Microbiology, 15(10):579–590, 2017.

11. A. Fritz, P. Hofmann, S. Majda, E. Dahms, J. Dröge, J. Fiedler, T. R. Lesker, P. Belmann, M. Z. DeMaere, A. E. Darling, et al. Camisim: simulating metagenomes and microbial communities. Microbiome, 7(1):1–12, 2019.

12. S. Garg. Computational methods for chromosome-scale haplotype reconstruction. Genome biology, 22(1):1–24, 2021.

13. I. Gregor, A. Schönhuth, and A. C. McHardy. Snowball: strain aware gene assembly of metagenomes. Bioinformatics, 32(17):i649–i657, 2016.

14. A. Gurevich, V. Saveliev, N. Vyahhi, and G. Tesler. Quast: quality assessment tool for genome assemblies. Bioinformatics, 29(8):1072–1075, 2013.

15. S. Hudault, J. Guignot, and A. L. Servin. Escherichia coli strains colonising the gastrointestinal tract protect germfree mice againstsalmonella typhimuriuminfection. Gut, 49(1):47–55, 2001.

16. H. Karch, P. I. Tarr, and M. Bielaszewska. Enterohaemorrhagic escherichia coli in human medicine. International Journal of Medical Microbiology, 295(6-7):405–418, 2005.

17. S. Koren, B. P. Walenz, K. Berlin, J. R. Miller, N. H. Bergman, and A. M. Phillippy. Canu: scalable and accurate long-read assembly via adaptive k-mer weighting and repeat separation. Genome research, 27(5):722–736, 2017.

18. D. Li, C.-M. Liu, R. Luo, K. Sadakane, and T.-W. Lam. Megahit: an ultra-fast single-node solution for large and complex metagenomics assembly via succinct de bruijn graph. Bioinformatics, 31(10):1674–1676, 2015.

19. H. Li. Minimap2: pairwise alignment for nucleotide sequences. Bioinformatics, 34(18):3094–3100, 2018.

20. Z. Li, Y. Chen, D. Mu, J. Yuan, Y. Shi, H. Zhang, J. Gan, N. Li, X. Hu, B. Liu, et al. Comparison of the two major classes of assembly algorithms: overlap–layout–consensus and de-bruijn-graph. Briefings in functional genomics, 11(1):25–37, 2012.

21. L. L. Ling, T. Schneider, A. J. Peoples, A. L. Spoering, I. Engels, B. P. Conlon, A. Mueller, T. F. Schäberle, D. E. Hughes, S. Epstein, et al. A new antibiotic kills pathogens without detectable resistance. Nature, 517(7535):455–459, 2015.

22. C. Luo, R. Knight, H. Siljander, M. Knip, R. J. Xavier, and D. Gevers. Constrains identifies microbial strains in metagenomic datasets. Nature biotechnology, 33(10):1045–1052, 2015.

23. B. A. Methé, K. E. Nelson, M. Pop, H. H. Creasy, M. G. Giglio, C. Huttenhower, D. Gevers, J. F. Petrosino, S. Abubucker, J. H. Badger, et al. A framework for human microbiome research. nature, 486(7402):215, 2012.

24. A. L. Mitchell, M. Scheremetjew, H. Denise, S. Potter, A. Tarkowska, M. Qureshi, G. A. Salazar, S. Pesseat, M. A. Boland, F. M. I. Hunter, et al. Ebi metagenomics in 2017: enriching the analysis of microbial communities, from sequence reads to assemblies. Nucleic acids research, 46(D1):D726–D735, 2018.

25. M. A. Moran. The global ocean microbiome. Science, 350(6266), 2015.

26. S. M. Nicholls, W. Aubrey, K. De Grave, L. Schietgat, C. J. Creevey, and A. Clare. On the complexity of haplotyping a microbial community. Bioinformatics, 37(10):1360–1366, 2021.

27. J. N. Nissen, J. Johansen, R. L. Allesøe, C. K. Sønderby, J. J. A. Armenteros, C. H. Grønbech, L. J. Jensen, H. B. Nielsen, T. N. Petersen, O. Winther, et al. Improved metagenome binning and assembly using deep variational autoencoders. Nature biotechnology, 39(5):555–560, 2021.

28. Y. Peng, H. C. Leung, S.-M. Yiu, and F. Y. Chin. Idba-ud: a de novo assembler for single-cell and metagenomic sequencing data with highly uneven depth. Bioinformatics, 28(11):1420–1428, 2012.

29. C. Quince, T. O. Delmont, S. Raguideau, J. Alneberg, A. E. Darling, G. Collins, and A. M. Eren. Desman: a new tool for de novo extraction of strains from metagenomes. Genome biology, 18(1):1–22, 2017.

30. R. Rizzi, S. Beretta, M. Patterson, Y. Pirola, M. Previtali, G. Della Vedova, and P. Bonizzoni. Overlap graphs and de bruijn graphs: data structures for de novo genome assembly in the big data era. Quantitative Biology, 7(4):278–292, 2019.

31. V. Sevim, J. Lee, R. Egan, A. Clum, H. Hundley, J. Lee, R. C. Everroad, A. M. Detweiler, B. M. Bebout, J. Pett-Ridge, et al. Shotgun metagenome data of a defined mock community using oxford nanopore, pacbio and illumina technologies. Scientific data, 6(1):1–9, 2019.

32. C. S. Smillie, J. Sauk, D. Gevers, J. Friedman, J. Sung, I. Youngster, E. L. Hohmann, C. Staley, A. Khoruts, M. J. Sadowsky, et al. Strain tracking reveals the determinants of bacterial engraftment in the human gut following fecal microbiota transplantation. Cell host & microbe, 23(2):229–240, 2018.

33. V. Somerville, S. Lutz, M. Schmid, D. Frei, A. Moser, S. Irmler, J. E. Frey, and C. H. Ahrens. Long-read based de novo assembly of low-complexity metagenome samples results in finished genomes and reveals insights into strain diversity and an active phage system. BMC microbiology, 19(1):1–18, 2019.

34. C. J. Stocks, M.-D. Phan, M. E. Achard, N. T. K. Nhu, N. D. Condon, J. A. Gawthorne, A. W. Lo, K. M. Peters, A. G. McEwan, R. Kapetanovic, et al. Uropathogenic escherichia coli employs both evasion and resistance to subvert innate immune-mediated zinc toxicity for dissemination. Proceedings of the National Academy of Sciences, 116(13):6341–6350, 2019.

35. A. Strazzulli, S. Fusco, B. Cobucci-Ponzano, M. Moracci, and P. Contursi. Metagenomics of microbial and viral life in terrestrial geothermal environments. Reviews in Environmental Science and Bio/Technology, 16(3):425–454, 2017.

36. K. Suvarna, D. Stevenson, R. Meganathan, and M. Hudspeth. Menaquinone (vitamin k2) biosynthesis: Localization and characterization of the mena gene fromescherichia coli. Journal of bacteriology, 180(10):2782–2787, 1998.

37. O. Tenaillon, D. Skurnik, B. Picard, and E. Denamur. The population genetics of commensal escherichia coli. Nature Reviews Microbiology, 8(3):207–217, 2010.

38. R. Vicedomini, C. Quince, A. E. Darling, and R. Chikhi. Strainberry: automated strain separation in low-complexity metagenomes using long reads. Nature Communications, 12(1):1–14, 2021.

39. M. Wu and J. A. Eisen. A simple, fast, and accurate method of phylogenomic inference. Genome biology, 9(10):1–11, 2008.

40. S. Zheng, S. Shao, Z. Qiao, X. Chen, C. Piao, Y. Yu, F. Gao, J. Zhang, and J. Du. Clinical parameters and gut microbiome changes before and after surgery in thoracic aortic dissection in patients with gastrointestinal complications. Scientific reports, 7(1):1–11, 2017.

